# An atlas of thyroid hormone receptors target genes in mouse tissues

**DOI:** 10.1101/2022.08.16.504127

**Authors:** Yanis Zekri, Romain Guyot, Frédéric Flamant

**Affiliations:** Institut de Génomique Fonctionnelle de Lyon (IGFL), CNRS UMR 5242, INRAE USC 1370, École Normale Supérieure de Lyon, 46 allée d’Italie, 69364 Lyon cedex 07, France

**Keywords:** thyroid hormone receptors, RNA-seq, ChIP-seq, atlas, target genes

## Abstract

We gathered in a single database available RNA-seq and ChIP-seq data to better characterize the target genes of the thyroid hormone receptors in several cell types. This database can serve as a resource to analyze the mode of action of the hormone. Also, it is an easy-handling convenient tool to obtain information on specific genes in regards to T3 regulation, or extract larger list of genes of interest based on the users’ criteria. Overall, this atlas is a unique compilation of recent sequencing data focusing on thyroid hormones, their receptors, mode of action, targets and roles which may profit researchers within the field. A preliminary analysis indicates extensive variations in the repertoire of target genes which transcription is upregulated by chromatin-bound nuclear receptor. Although it has a major influence, chromatin accessibility is not the only parameter that determines the cellular selectivity of hormonal response.

## Introduction

Thyroid hormone (3,3’, 5-triiodo-L-thyronine or T3) exerts a broad influence on vertebrate development and adult physiology. It is notably known to trigger frog and fish metamorphosis (Tata, 2006). During mammalian development, it is required for proper brain maturation and bone growth and becomes a main regulator of energy metabolism in adults. At the cellular level, T3 often stimulates differentiation (Gandrillon et al., 1994; Robson et al., 2000; Billon et al., 2001; Boukhtouche et al., 2010) and mitochondrial metabolism (Wrutniak-Cabello et al., 2001).

T3 exerts its influence by binding to nuclear receptors TRα1, TRβ1 and TRβ2 (collectively TR) encoded by the two *Thra* and *Thrb* genes (Flamant, 2016). TR act as heterodimers with other nuclear receptors, RXRs. They bind to specific Thyroid Response Elements (TRE) constituted by doublets of the AGGTCA half-site (DR4 elements) (Velasco et al., 2007) present in the regulatory sequences of genes, which transcription is upregulated upon T3 binding (Yen, 2001). Although both *Thra* and *Thrb* genes have distinct expression patterns, all cell types express at least one of the two genes, and are thus potentially T3 responsive. However, it seems that T3 activates the transcription of different genes across cell types, which explains why it exerts different physiological influences depending on the tissue. The molecular basis for the cell-type specificity of T3 response is currently unclear.

Because of the broad influence of T3 on gene expression in many cell types, an inventory of the TR target genes should enlighten many important developmental and physiological functions. This represents however a tremendous task. A previous compilation of genome wide expression analyses, mainly based on microarray analyses, identified only few genes, which were regulated by T3 in several cell types or tissues (Chatonnet et al., 2015). Since this early attempt, a number of novel studies have been performed in mice. Both RNA-seq and ChIP-Seq analyses have been used to better characterize TR target genes in several cell types or tissues. We present here an atlas of the currently available datasets, which can be used as a novel resource to explore the many function of T3 and from which we try to extract general conclusions on T3 signaling.

## Methods

### Transcriptome analysis

Raw data were downloaded from Gene Expression Omnibus. Their accession numbers are listed in Table 1. Sequence reads were aligned on the GRCm38 (mm10) reference genome using Bowtie with default setup (Galaxy Version 2.2.6.2). Reads with an alignment quality inferior to 10 were eliminated. The number of reads assigned to each gene was calculated using htseqcount (Galaxy Version 0.6.1galaxy3). Finally, differential analysis of gene expression was performed with DESeq2 (R package, Version 1.34.0). Differentially expressed genes (DEG) were defined as genes with a mean average expression > 10 and an adjusted p-value <0.05. No threshold was used for log2 fold-change. Using these parameters, genes upregulated were marked with U, downregulated with D, not sensitive to T3 with N and genes not expressed with NA. For time course analyses of T3 response, each timepoint was compared to the control samples, prepared from either hypothyroid or euthyroid mice. Importantly, only genes that were regulated in at least one tissue were kept in the atlas. Gene ontology analysis was made using the Gene Ontology Resource (http://geneontology.org).

**Table 1:**
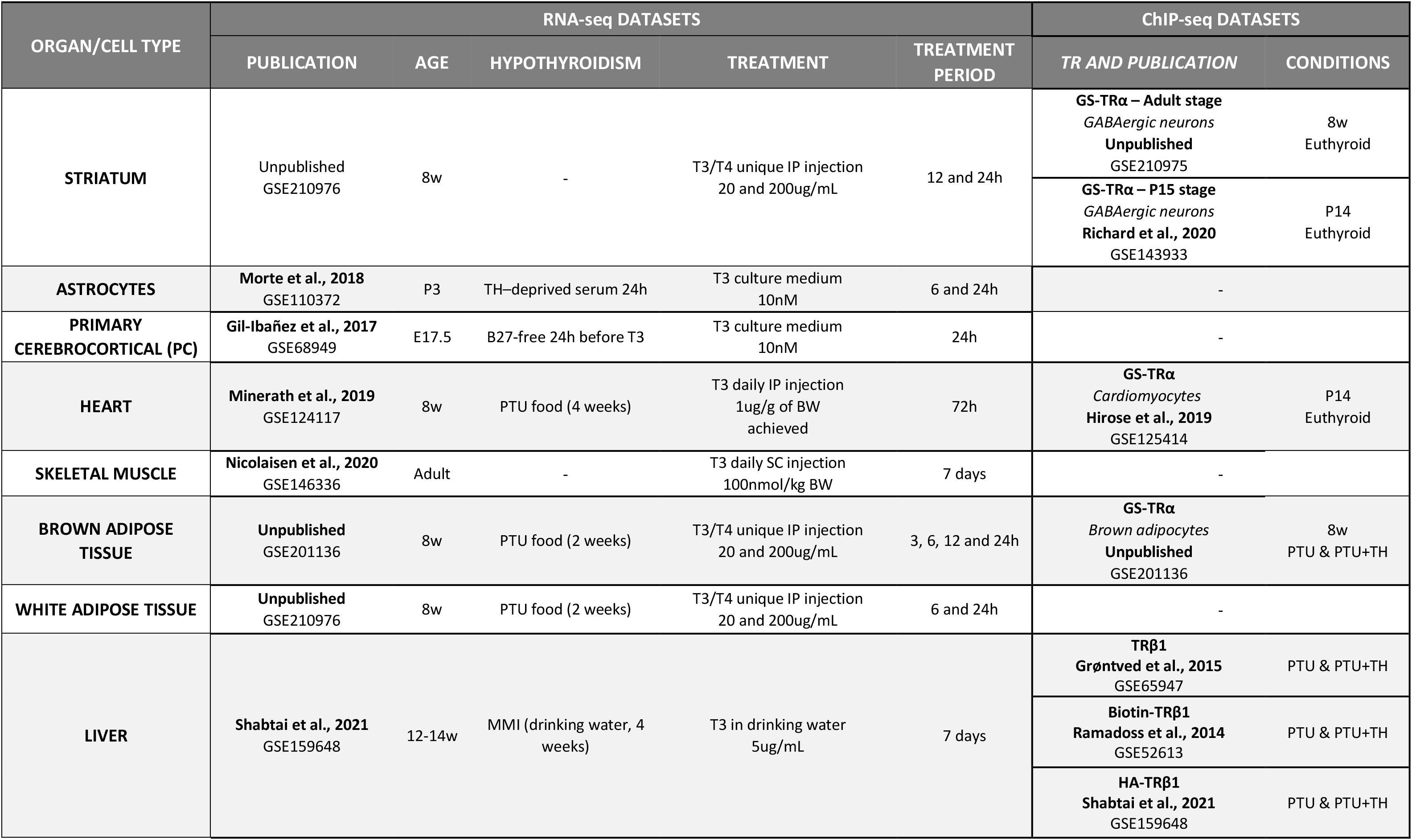
Data included in the atlas

### Cistrome analysis

Raw data was downloaded from Gene Expression Omnibus. Their accession numbers are listed in Table 1. Sequence reads were aligned on the GRCm38 (mm10) reference genome using Bowtie with default setup (Galaxy Version 2.2.6.2). MACS (Galaxy Version 2.1.1.20160309.0) was used for peak calling and peaks with a score inferior to 60 were filtered out. Genes within 30kb of peaks were called out using GREAT (http://great.stanford.edu/public/html/). According to our estimations (Fig. 2B), this distance maximizes the ratio of T3-responsive genes, without excluding genes which have been well characterized as TRα1 target genes, such as *Klf9* or *Hr* (Gil-Ibañez et al., 2013). Bigwig files were obtained by converting the Bedgraph files from MACS2 with the Wig/BedGraph-to-bigWig converter (Galaxy Version 1.1.1). Peaks were visualized by uploading Bigwig files on the Integrative Genome Viewer (2.12.3). The distribution of distances of TRBSs around TSSs, as well as the distribution of TRBSs in the genome, were assessed using PAVIS (https://manticore.niehs.nih.gov/pavis2/).

**Figure 1:**
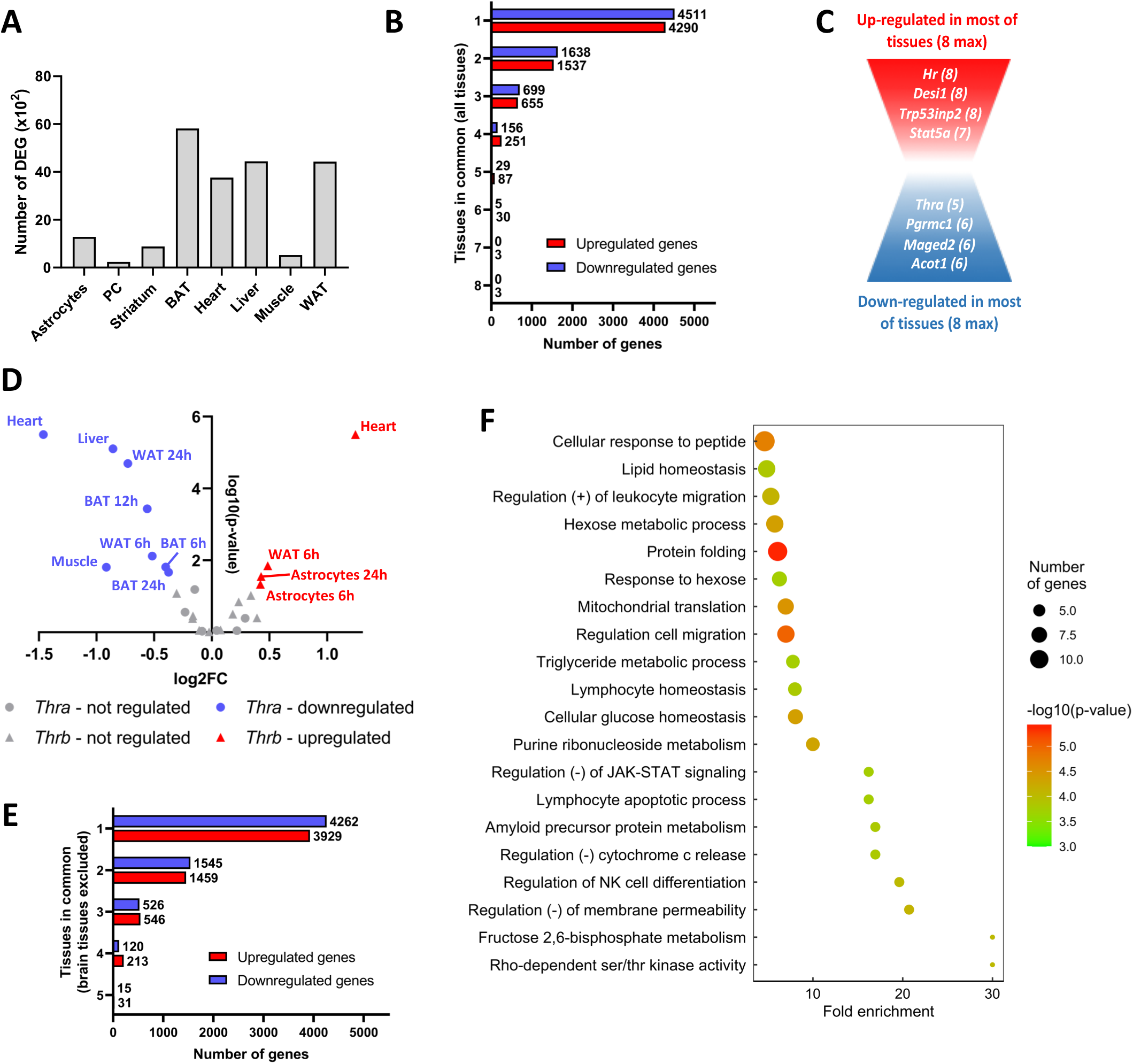
Thyroid hormone regulate the expression of tissue-specific genes. **(A)** Number of differentially expressed genes (DEG, x10^2^) in the available tissues. **(B)** Number of shared genes up or downregulated among tissues. Genes were classified in 8 categories, depending on the number of tissues within which they were regulated. **(C)** Schematic representation of the up and downregulated shared genes among tissue. **(D)** Regulation of *Thra* (circle) and *Thrb* (triangle) by thyroid hormones. Log_2_ fold-change and statistical significance were plotted for each timepoint of each tissue. When the regulation was significant (i.e, when −log_10_ p-value > 1.3, meaning p-value < 0.05), tissues within which *Thra/Thrb* were up or downregulated were colored in red and blue respectively. Tissue name and timepoint of the kinetics were attributed to each point. **(E)** Number of genes up or downregulated among tissues, excluding brain tissues. Genes were classified in 5 categories, depending on the number of tissues within which they were regulated. **(F)** Gene ontology dot plot of the genes upregulated in at least 4 of the 5 non-brain tissues. Some of the terms were shortened to increase readability without affecting the meaning.

**Figure 2:**
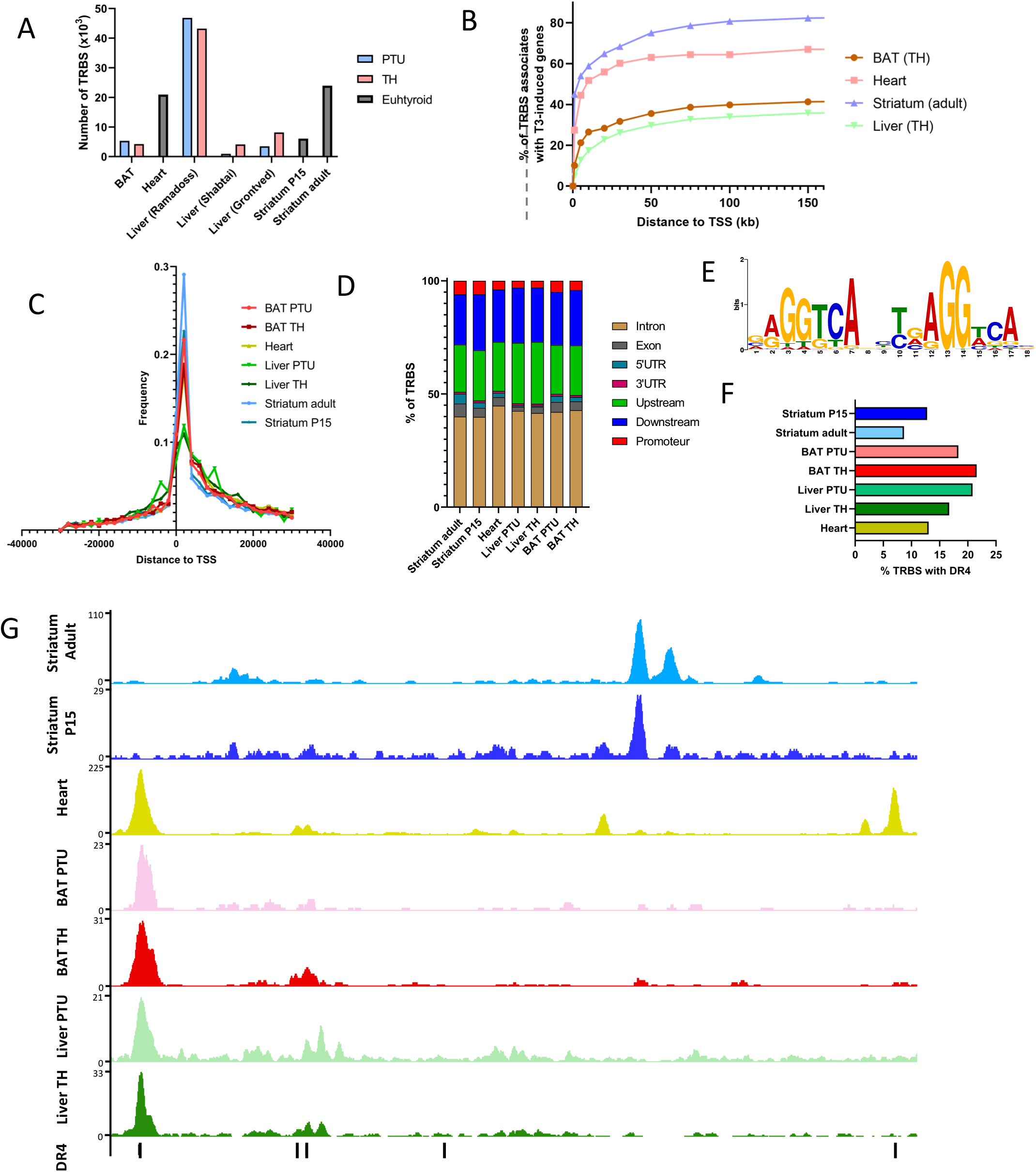
TR chromatin occupancy. **(A)** Number of TR binding sites (TRBS) in each of the available tissue. The different thyroid status are indicated by a color code (blue: hypothyroid by PTU exposure, pink: TH injection, grey: euthyroid). **(B)** Percentage of genes upregulated by T3 that possess a TRBS depending on the chosen distance (in kilobase) to attribute a TRBS to a gene. We generally observe 3 phases: a very high-enrichment of T3-induced when considering TRBS within 10kb, a high-enrichment for TRBS within 30kb and a plateau above 50kb. Thus, we chose 30kb as a distance to attribute a TRBS to a gene, as a compromise to maximize the number of TR target genes while minimizing false-positives. **(C)** Frequency of TRBS relatively to their distance of the TSS. **(D)** Distribution of TRBS in different gene regions. **(E)** Consensus sequence as identified by *de novo* motif search in brown adipocytes. All the available tissues display the similar DR4 element as the top-enriched motif. **(F)** Percentage of TRBS that possess a DR4. **(G)** Extract of the Integrative Genome Viewer (IGV) in the promoting region of *Hcn2.* Each tissue is represented by a different color and DR4 elements are represented by black vertical lines at the bottom of the IGV shot.

*De novo* motif search was made by submitting filtered peaks to the MEME-ChIP program (MEME Suite v5.4.1 https://meme-suite.org/meme/) (Machanick and Bailey, 2011) looking for motifs between 6 and 20 nucleotides. Only motifs with a p-value < 0.05 were conserved. To estimate the proportion of TRBS that possess a DR4, we crossed our datasets with a BED file that gathers the more than 70 000 DR4 present in the genome (Table S1). This table was built using FIMO from the normalized probability matrix produced by MEME-ChIP (Grant et al., 2011).

## Results

### Definition of TR target genes

T3 alters gene expression in exposed cells both directly, by binding the chromatin bound TR, and indirectly, because TR target genes can encode transcription regulators, which rapidly generate a secondary response. Unraveling these two overlapping processes is difficult, although this can be done *in vitro* with a translation inhibitor (Gil-Ibáñez et al., 2014; Gil-Ibañez et al., 2017; Morte et al., 2018). We will use here two simple criteria to define TR target genes:

1. Genes which are up-regulated after T3 treatment of hypothyroid mice. The initial T3 depletion is important, because some genes are more sensitive to it than to an excess of T3 (Yen et al., 2003). Hypothyroidism is most of the time obtained by pharmacological means with at least two weeks of treatment with either propyl-thio-uracyl (PTU) or methimazole (MMI). *In vitro*, serum used for cell cultures should be depleted of T3. When a time-course analysis is performed, a rapid response, within hours, is a good indication that the transcriptional activation is a direct consequence of TR-mediated regulation.
2. Regulatory sequences are occupied by either TRα1 or TRβ1/2, as evidenced by chromatin immunoprecipitation. Up to now, the difficulty to raise high quality antibodies against TR, which are not abundant proteins, has limited the production of ChIP-seq datasets to liver (Grøntved et al., 2015; Hönes et al., 2017). This technical limitation is now commonly circumvented by using tagged receptors, for which several transgenic mice have been produced (Minakhina et al., 2020; Shabtai et al., 2021). The availability of a “floxed” construct (Hirose et al., 2019) allows to address chromatin occupancy by TRα1 in a single cell type within a heterogeneous tissue.

### Presentation of the atlas

Table 1 depicts all the mouse datasets that are currently available in the literature to our knowledge, linking transcriptome (RNA-seq) and cistrome (ChIP-seq) data to facilitate the identification of TR target genes. We excluded data obtained on cell lines, and included data obtained from primary cell cultures. Overall, we collected 8 RNA-seq datasets accounting for different brain and non-brain tissues, covering cell types of very different functions and embryonic origin. It is worth noting that these datasets emerge from different protocols, using: (1) mice of different ages, (2) submitted or not to different hypothyroidism procedures and (3) treated with different period and doses of thyroid hormone, (4) administrated in different manners. These differences undoubtedly trigger responses with very various magnitudes, making difficult the quantitative comparisons between tissues.

We also collected 7 ChIP-seq datasets, including two for GABAergic neurons at different developmental stages, and 3 for liver. Some of these datasets display different thyroid status allowing to study the consequences of T3 binding to TRs on chromatin occupancy.

To avoid any bias, we reanalyzed all the datasets using a homogeneous analytical pipeline for sequence mapping and counting of genes coverage. Further details are presented in the methods section. Finally, all these data were combined into a single atlas (Table S2), where one can find expression and chromatin-occupancy data for each gene. A tutorial has been added to the atlas in a dedicated worksheet to help the reader in its use.

### Differentially expressed genes

Differential gene expression analysis identified thousands of genes which expression level is modified, either *in vivo* or *in vitro*, after T3 treatment (Table S2). The number of differentially expressed genes (DEG) varies extensively from one cell population to the other (Fig. 1A). This suggests that different cell types display different sensitivity to T3. According the Table S2, this feature cannot simply be explained by variations in the expression of *Thra* and *Thrb.* Noteworthy, this difference may also account for the different protocols used. In particular, as mice were not made hypothyroid prior to T3 treatment, the effects on the muscle transcriptome are less visible.

The vast majority of DEG are not shared by the different cell types (Fig. 1B). To the well-known *Hr* gene (Thompson, 1996) which encodes the *Hairless* transcription cofactor, one can add *Desi1, Trp53ind* and *Stat5a* (Fig.1C). The presence of this last gene, which encodes a member of the STAT family of transcription factors, raises interesting possibilities for cross-talks between T3 signaling and other signaling pathways, which remain to be explored. While no genes were repressed by T3 in all cell types, we still identified a set of genes with this recurrent pattern. Interestingly, it included *Thra* itself, which might provide a molecular basis for a negative feedback loop. Thus, we looked more precisely at *Thra* and *Thrb* regulation and observed that *Thrb* had an opposite trend (Fig. 1D), although not as pronounced as in metamorphosing amphibians (Tata, 2000).

Because the response of cells outside of the brain is thought to be mainly metabolic, while neural cells T3 response is described as mainly relevant to terminal differentiation and maturation, we analyzed separately the 5 datasets excluding brain tissues. Although they shared a higher fraction of DEG (Fig. 1E), the overlap between the sets of DEG remained limited. We used gene ontology to identify biological functions enriched in these cell types and found a significant enrichment for several gene sets (Fig. 1F). While the category “lipid homeostasis” and “mitochondria translation” was expected, other items were more surprising, notably those suggesting an immune infiltration. Overall, this analysis indicates that different cell types display very different responses to T3, with a few similarities when considering only the responses of cell types outside of the brain.

### Chromatin occupancy by TR

The number of binding sites occupied by TR in the chromatin (TRBS) identified by ChIP-seq analysis is highly variable (Fig.2A), which might reflect either technical variations, or a genuine influence of the cellular environment on chromatin occupancy. The fact that 3 studies address TR occupancy in liver, in which T3 response is almost exclusively mediated by TRβ1, allows to identify the technical sources of this variability. The overexpression of tagged TRβ1 by an adenovirus vector (Ramadoss et al., 2014) greatly helps for the detection of chromatin occupancy sites, but probably facilitates binding of the receptor to chromatin. In particular, it erases the influence of T3 on chromatin accessibility, which is observed in the two other liver studies. Also, it drastically increases the mean number of peaks within the same gene. In the following, we will favor the study which uses transgenic mice with a tagged TRβ1 (Shabtai et al., 2021) because it takes advantage of the tagging of TRβ1 to produce high quality data without taking the risk of generating artificial occupancy by overexpression.

The common way to combine transcriptome and cistrome data is to set an arbitrary distance between the TRBS and the transcription start site (TSS). The gene is then called a TR target gene if its expression is T3 responsive and if a TRBS is present within the interval given by this threshold distance. Different studies vary by the definition of this distance, which results from a compromise: long distance will produce more false-positives, while a small distance will increase the rate of false-negatives. For our novel analysis, we calculated the fraction of T3 responsive genes that possess a TRBS depending on the distance choice (Fig. 2B). We concluded that a 30 kb distance was a good compromise. Intriguingly, the distribution of the TRBS with respect to the TSS is not symmetrical, and the fact that they are more frequent downstream to the TSS is at first site counterintuitive (Fig. 2C). Accordingly, a large fraction of the TRBS are found in introns (fig. 2D).

ChIP-seq does not only capture the direct association of proteins to DNA, but also the indirect connections mediated by protein-protein interactions, also called “protein tethering”. Motif analysis can be viewed as a way to better identify all the binding proteins and also to define consensus sequences for each protein binding to chromatin. We performed *de novo* motif search in each dataset and the results were remarkably similar. A single consensus sequence was detected which corresponds to the so-called DR4 element (fig. 2E). The association of TRα1/RXRα heterodimer on this element has been previously analyzed in great details by X-ray crystallography (Rastinejad et al., 1995). TR occupy the downstream half-site, while RXR bind the 5’ half-site. The sequence of the 4 nucleotides spacer is less precisely defined but the last 2 nucleotides also establish contact with TR. Depending on statistical threshold for similarity, the detection of the consensus sequence in TRBS varies. Using a file compiling all the DR4 in the mouse genome (Table S1), we found DR4 elements in only 12-25% of the identified TRBS (Fig. 2F), which leaves open the possibility for other mode of association of TR with chromatin.

Thus, DR4-like elements occupied by TR according to ChIP-seq analysis represent only few thousands out of 70 394 DR4-like elements present in the mouse genome. This outlines that chromatin accessibility is an important factor, which governs the occupancy of DR4 elements by TR. Accordingly, within the same genomic location, we can observe very different patterns (Fig. 2G). Some TRBS will be detected only in non-brain tissues, with the underlying presence of a DR4 site. Some will be unique, with or without the presence of a DR4, while some DR4 will be present in unoccupied sites.

### Evolution of chromatin occupancy during cell differentiation and after T3 treatment

The previous analysis outlines striking differences in chromatin occupancy among cell types. The available ChIP-seq datasets allow to better understand the source of these variations. In particular, the analysis of chromatin occupancy in the GABAergic neurons of striatum has been performed using the same protocol, at two different life stages: juvenile, when neuronal circuits are still immature, and adult. The number of TRBS is much higher in adults, which suggests a better accessibility of putative binding sites. This is not the only parameter which evolves with time however, because some TRBS are also lost during striatum maturation (Fig. 3A). The fact that the DR4 containing TRBS represent a smaller fraction among the TRBS gained at adult stage raises an interesting possibility: it might reflect the assembly of multiprotein complexes containing TRα1. These would contact chromatin at several different points and thus produce several TRBS. It is indeed frequently observed that T3-responsive genes often contain more than one TRBS, among which a single DR4 element is present.

**Figure 3:**
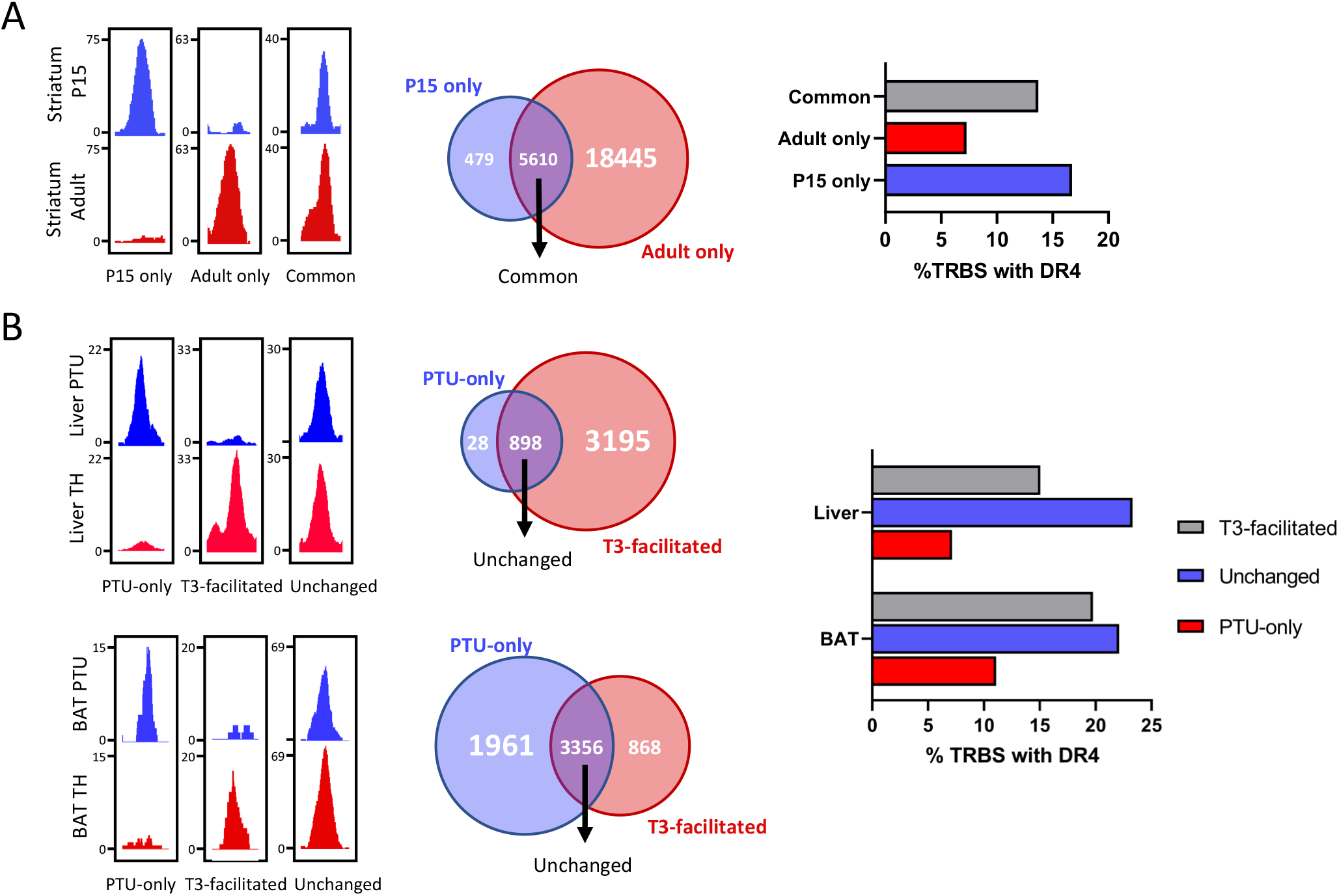
TR plasticity during development and T3 treatment. **(A)** *(Left)* Integrative Genome Viewer shots of TRBS categorized as only appearing in P15 striatum, adult striatum or present in both. *(Middle)* Venn diagram of TRBS present on each of the defined categories. *(Right)* Percentage of TRBS that possess a DR4 element. **(B)** *(Left)* Integrative Genome Viewer shots of TRBS categorized as only present in the PTU condition, facilitated by T3 or present independently of T3 status. *(Middle)* Venn diagram of TRBS present on each of the defined categories. *(Right)* Percentage of TRBS that possess a DR4 element. This analysis was made for liver *(top)* and brown adipocytes *(bottom)*.

We also address the influence of T3 on chromatin occupancy, a question which was previously investigated in depth in liver studies (Grøntved et al., 2015; Shabtai et al., 2021). In liver, T3 increases the number of TRBS, and this is not due to an induction of *Thrb* expression. Again, some TRBS are lost while others are gained soon after T3 treatment (Fig. 3B). The enlargement of the cistrome after T3 treatment is unlikely to become a general rule, as an opposite trend is found in brown adipocytes, although this conclusion relies on a single dataset.

### Combination of transcriptome and cistrome data

RNA-seq indicates that an equivalent number of genes are downregulated or upregulated after T3 treatment (Fig. 4A), which might lead to question the idea that liganded TR are only transcription activators. However, 45% of genes upregulated in the BAT after 3h of T3 possess a TRBS within 30kb of the TSS, a proportion that fell to 35% overtime, certainly due to the accumulation of indirect regulations. However, this was the case for less than 10% of downregulated genes, a proportion similar to genes insensitive to T3 (Fig. 4B). This suggests that the negative regulation of gene expression is not directly mediated by liganded TR and could be secondary for example to the upregulation of genes encoding transcription repressors.

**Figure 4:**
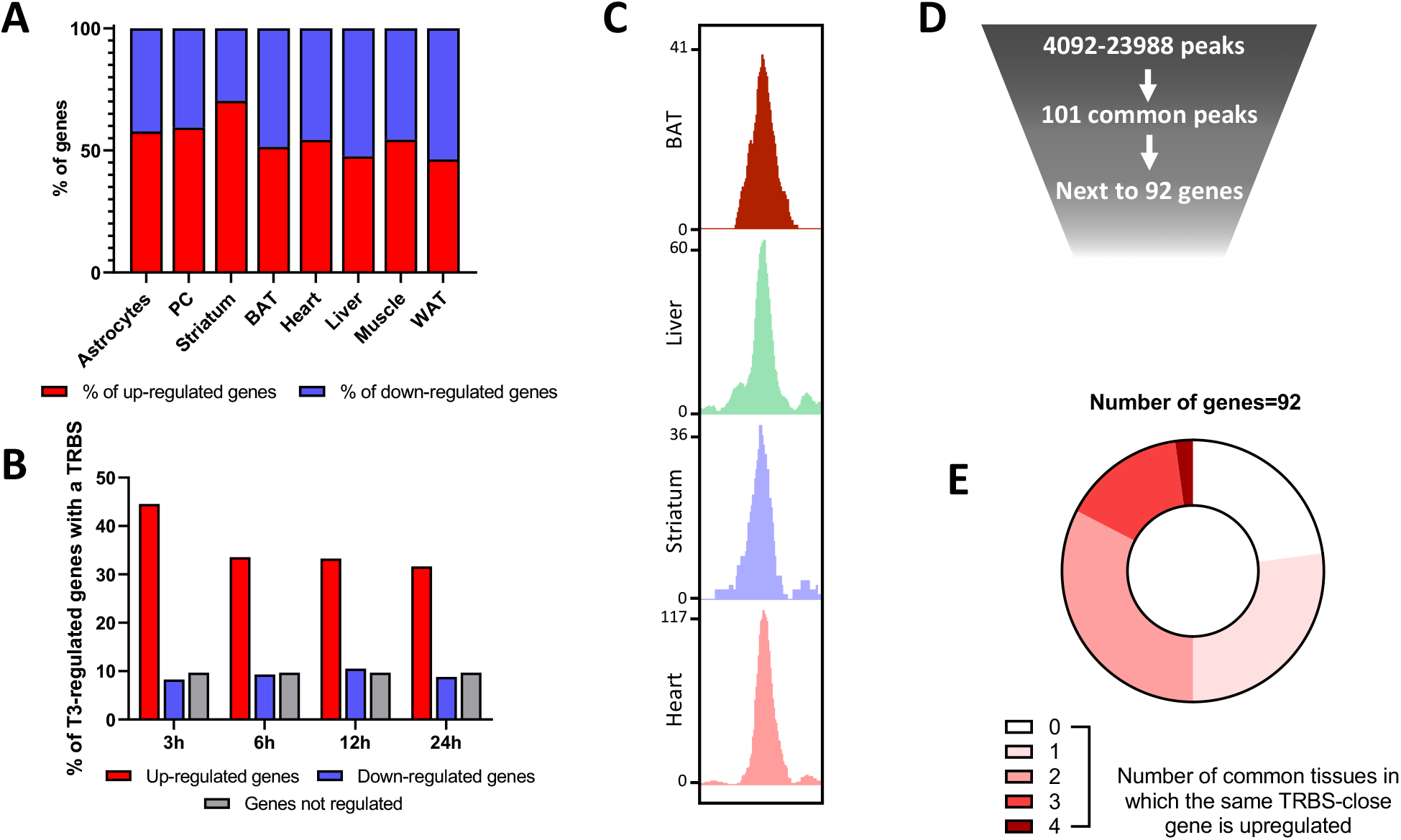
Combined analysis of RNA-seq and ChIP-seq data gives insights on TR mechanisms of action. **(A)** Percentage of genes up or downregulated by thyroid hormone in the different available tissues. **(B)** Percentage of genes which possess a TRBS within 30kb of their TSS among genes whose expression in the BAT is regulated or not by T3 (upregulated in red, downregulated in blue and a set of randomly-picked not regulated genes in grey). **(C)** Integrative Genome Viewer shot of a TRBS shared in the four different available tissues. **(D)** Schematic representation of the rarity of common TRBS. **(E)** Circle diagram of T3 sensitivity for the 92 genes that are close to the 101 common TRBS. These genes are divided in 5 categories, depending in how many of the 5 tissues their expression is induced by T3.

A large fraction of the TRBS were not located next to T3 responsive genes which can receive three non-exclusive explanations. (1) First, it might result from chromatin folding which enables TR to act at very long distance, as other nuclear receptors do (Le Dily and Beato, 2018). (2) Another possibility would be that a large fraction of TRBS are not functional, reflecting a mode of association of TR with DNA which does not allow the recruitment of transcription coactivators. This could eventually result from the labile interaction of TR with low-affinity DNA binding motives, as recently suggested by *in vitro* assays (Flamant et al., 2022). (3) Finally, the cellular context might be determinant, and cell-specific coactivators might be involved in converting T3 binding to chromatin associated TR into transcription activation. To better consider this last possibility we focused the analysis on the TRBS shared in the 4 available tissues (Fig. 4C). These 101 shared TRBS (Fig.4D) are located next to 92 genes and 40% of them contain DR4 elements, a ratio which is above the average frequency. However only 2 of these genes are T3 responsive in the analyzed cell types (Fig. 4E). This evidences that TR binding is necessary but not sufficient for T3 transactivation, and that cell-specific parameters are involved in converting TR binding into transcriptional activation.

## Discussion

The present study addresses the mechanisms of T3 mediated-transactivation by gathering and analyzing in a uniform manner the currently available datasets. This survey outlines that different cell types respond to T3 stimulation in a very different manner. The overlap between the repertoires of TR target genes in different cell-types remains limited, even for cell-types in which T3 activates energy metabolism and mitochondrial functions, like cardiomyocytes, hepatocytes and adipocytes.

We can speculate that this variability occurs also within each tissue. Indeed, the T3 response analyzed at whole organ level results from the combination of the T3 response of several cell types, which might generate confusion in data interpretation. For example, adipocytes are only 50% of the cells found in white adipose tissue and their high heterogeneity just starts to be perceived (Emont et al., 2022). The use of single-cell methods for T3 response could give insights on which target genes is regulated by which cell-subtypes, as already seen with other hormones (Hoffman et al., 2020).

Chromatin occupancy is a crucial criterion in our definition of TR target genes to eliminate genes most likely regulated indirectly. Obviously, classical chromatin-immunoprecipitation only provides a one-dimensional picture for binding sites, forcing us to choose an interval distance within which we can attribute a T3-sensitive gene to a TRBS. Despite our efforts to estimate the best compromise, we may have missed active TRBS due to this limitation. The growing use of methods considering the two-dimensional conformation of chromatin may overcome these problematics (Belton et al., 2012; van de Werken et al., 2012; Han et al., 2018).

Previous *in vitro* studies have identified non-DR4 consensus sequences able to mediate T3/TR/RXR binding and transactivation, called ER6 (everted repeats with a 6 nucleotides spacer) and IR0 (inverted repeats half-site without spacer) (Paquette et al., 2014). This was previously denied by scattered studies (Chatonnet et al., 2013; Richard et al., 2020). Here, the combined analyses of multiple datasets identified DR4 as the unique consensus motif significantly enriched which raises some doubts on the physiological relevance of the non-DR4 binding sites for T3 mediated transactivation.

Overall this global analysis outlines a basic problem which should be addressed in further research: the genome contains more than 70 000 DR4-like sequences, which are putative TRBS. Among these, only few thousands are actually occupied by TR, according to ChIP-seq analysis. Thousands of genes are proximal to these TRBS, but the transcription of only few hundreds of these is upregulated by T3. This suggests that only a fraction of chromatin bound TR convert T3 binding into transcriptional activation. In the following, we consider promising approaches to overcome the current limitations of the RNA-seq/ChIP-seq which should allow to tackle this question.

(1) Many TR target genes encode for extracellular proteins: growth factors, neurotrophines, etc. Therefore, in a tissue, the cellular response to these factors is difficult to separate from a direct, cell-autonomous response to T3. This has been notably exemplified in brain where interaction between neighboring cell types is a common theme (Fauquier et al., 2014; Richard et al., 2020). In mice the availability of *Thra* and *Thrb* “floxed” alleles enables to address this point. TR target genes are expected to be downregulated when the T3 response is selectively blocked in the considered cell type. This provides an additional and useful criterion to define TR target genes (Flamant et al., 2017; Richard et al., 2020).

(2) Cross-species comparisons among mammals or vertebrates could outline the gene regulation that are the most relevant for conserved function of T3. Only few attempts have been made in that direction, but the regulation of *Klf9* by TR is clearly conserved between mammals and amphibians (Bagamasbad et al., 2015) and pivotal for neuronal maturation (Avci et al., 2012).

(3) Other genomic assays are required to address the influence of chromatin remodeling on cistrome and define the consequences of TR binding, notably on histone tail modifications (Præstholm et al., 2020). This should help to address the possibility that many “silent TRBS” exist.

(4) The development of high-throughput functional assays to demonstrate the capacity of TRBS to mediate T3 transactivation is needed. A proof of principle has been recently obtained that the so-called synthetic STARR-seq approach can address this question *in vitro* (Flamant et al., 2022).

In conclusion, we gathered in a single database the sequencing data of T3 response and TR occupancy in several tissues. This collection sheds the light and opens the discussion on several mechanistic aspects of TR signaling including TR-mediated repression, hormonal-dependent chromatin occupancy or the nature of TRE. More importantly, this atlas represents a highly valuable tool that could be useful for the entire thyroid hormone community. It can be used to quickly obtain information on specific genes in regards to T3 regulation or TR binding, or extract larger list of genes of interest. For example, the list of direct TR target genes in neurons was recently used to screen the effects of around 300 chemicals on thyroid signaling in order to identify and test *in vitro* the most active compounds *in silico* (Zekri et al., 2021). Thus, this atlas is a quickly accessible, easy handling resource to study the thyroid hormone target genes, their function and potential disruption in mice.

## Acknowledgments

We thank Catherine Cerrutti and Gerard Benoit for the advices about sequencing analysis as well as Benjamin Gillet, Sandrine Hughes and the PSI platform of the IGFL for deep sequencing. We also thank Emmanuel Quemener from Center Blaise Pascal/ENSL for the development and maintenance of the ENS Galaxy portal with the help of SIDUS (Single Instance Distributing Universal System). This work was supported by the European Union’s Horizon 2020 research and innovation program under grant agreement no. 825753 (ERGO).

## Supplementary figures

**Table S1: BED file of DR4 elements in the mouse genome**

**Table S2: Atlas of TR target genes**

